# Self-assembled cell-scale containers made from DNA origami membranes

**DOI:** 10.1101/2024.02.09.579479

**Authors:** Christoph Karfusehr, Markus Eder, Friedrich C. Simmel

**Author notes:** Contributing authors.

## Abstract

DNA origami provides a methodology for the sequence-programmable generation of precisely defined molecular nanostructures with sizes of order 100 nm. A new frontier for the field is the generation of superstructures made from DNA origami subunits, which requires other self-assembly strategies than those used for DNA origami itself. Challenges faced by current approaches include the increasing complexity, cost and development time for the structures and off-target assembly. Here, we demonstrate how radially symmetric origami subunits that are inspired by the structure and interactions of lipids organize into giant DNA origami monolayer membranes that can be readily programmed to form vesicles or hollow tubes with diameters ranging from 100 nm to over 1 µm. DNA origami membranes are an unprecedented approach for compartmentalization that opens up new possibilities for bottom-up biology and cell-scale soft robotics.

---

Compartmentalization is one of the hallmarks of living systems and is utilized across many length scales in biology. It allows the creation and control of local environments, in which chemical processes are efficiently performed and separated from their surroundings. Compartments often define modules that group functionally related components, which are optimized to perform distinct and specific tasks, facilitating the division of labor among different modules. They also enable the hierarchical organization of living matter into higher order structures, which ultimately gives rise to the complexity and diversity of life-forms observed on Earth.

At the molecular level, compartmentalization is often achieved by encapsulation within hollow containers, which are typically either crystalline cages formed by their monomeric constituents or self-organized membranes without long-range order (Figure 1).

**Fig. 1.**
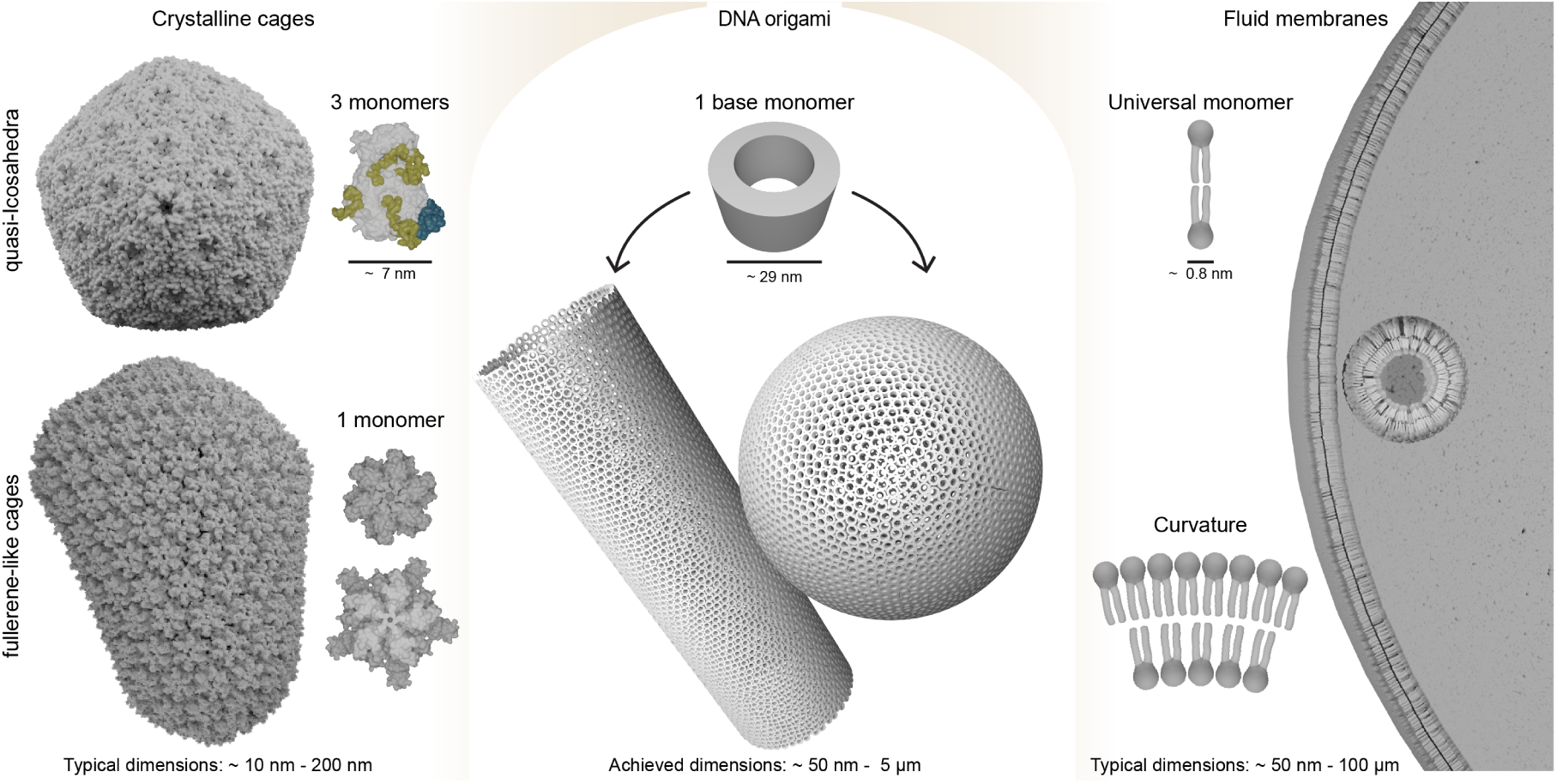
DNA origami (‘Dipid’) membranes combine the construction principles of protein cages and lipid bilayer membranes. In the left panel, rigid protein assemblies such as the carboxysome (PDB: 8B12) or a fullerene-like, mature HIV capsid (3J3Q, pentamer: 8CKW, hexamer: 8CKV) [5, 25, 26] are shown. The carboxysome is constructed from an asymmetric unit, composed of three distinct protein types (gray, green and blue). By contrast, the HIV capsid assembles from a single monomer unit that can arrange into pentameric and hexameric sub-assemblies. The right panel depicts the assembly of lipid bilayer membranes from amphiphilic lipids. Isotropic and weak interactions between the lipid monomers result in fluid and flexible membranes, whose curvature and overall size can be controlled by the shape of the lipids. DNA origami membranes (middle panel) are based on semi-, barrel-like base monomers with approximately isotropic nearest neighbor interactions. Similar as for lipids, assemblies with different curvatures can be realized with different versions of the base monomer. Protein structures were visualized in Blender, using the MolecularNodes plugin [27].

Biological examples of cage-like containers are the protein-based bacterial organelles such as the carboxysome or ferritin, or the capsids of viruses. Both types of containers are often found to display quasi-icosahedral symmetry, in which the subunits locally arrange in hexagonal or pentagonal symmetry. Regardless of the number of hexagonal subunits present, there are always precisely twelve pentameric subunits, a fundamental characteristic, which is utilized in the well-known Caspar-Klug (CK) construction of viral capsids [1]. Not all viruses faithfully follow the CK rules, however [2]. For instance, the HIV capsid is assembled from only a single type of capsid protein, which can form both pentameric and hexameric subunits [3]. The resulting cone-shaped capsid deviates from the standard quasi-icosahedral shape, forming fullerene-like cones, still obeying the 12 pentamer rule [4, 5].

On the scale of hundreds of nanometers and above, biology utilizes less ordered and more dynamic assemblies for compartmentalization, in particular lipid bilayer membranes and liquid-liquid phase separation [6]. Lipid membranes are formed by amphiphilic lipids, which compared to capsid proteins are rather flexible monomeric subunits that are held together by the hydrophobic effect and weak monomer-monomer interactions. Curved membranes are created by varying the aspect ratio between the sizes of the lipid head and tail groups. In contrast to rigid protein cages, lipid bilayer membranes, vesicles, and tubes can self-assemble from a single monomer type at minimal energetic input, and with considerable defect tolerance.

With applications in nanotechnology and synthetic biology in mind, a wide range of efforts have recently focused on designing and utilizing synthetically generated containers for compartmentalization. Among others [7–11], this involved the engineering of naturally occurring [12] or de novo designed protein cage formers [13, 14] and the use of amphiphilic molecules for the realization of vesicle-inspired peptidosomes [15], polymersomes [16], or DNAsomes [17]. Using the DNA origami technique, researchers have created capsid-mimicking DNA cages of sizes between 90 and 300 nm that were self-assembled from small numbers of rigid origami monomers [18, 19]. Among the benefits of using DNA origami for such containers is the straightforward chemical modification of the subunits, which, in principle, allows the sequence-addressable functionalization of the containers with nanoscale precision.

Until now, the creation of DNA origami superassemblies has utilized design paradigms that require precise control over subunit shapes and interactions [20, 21]. This approach is ideal if the realization of supramolecular containers with an exact number of monomers and geometric dimensions is desired [18, 19, 22]. However, this method presents challenges when scaling up to larger compartments. Creation of large compartments from unique subunits is prohibitive in terms of material use, monomer design effort, and assembly times [23, 24]. By definition, rigid designs are not adaptive and cannot be easily modified to other shapes or geometric dimensions.

In the present work, we sought to combine the nanoscale precision enabled by DNA origami design with the more flexible architecture of membrane-like assemblies. To this end, we developed an approach based on DNA origami subunits termed Dipids, whose architecture and interactions are inspired by the structure of lipids. Dipids are designed as isotropic sticky discs [2] that bind to their neighbors via a network of flexible DNA strands with weak hybridization interactions. We show that Dipids enable the assembly of diverse membrane objects like vesicles and tubes, ranging in size from the nanometer to the micron scale. Remarkably, all structures self-assemble from a single, rationally designed Dipid monomer. Dipid variants with different shapes and binding strengths can be easily derived from a single, easily modifiable DNA origami base unit.

## Design principles for lipid inspired origami tectons

Our starting point is the observation that lipid-lipid interactions within a membrane are largely isotropic. We therefore sought to construct a monomer without strong directional bonding, ideally a barrel-like structure with cylindrical symmetry. Among other DNA-based approaches [29, 30], DNA origami [31, 32] is an established self-assembly technique that allows the creation of almost arbitrarily shaped molecular objects from DNA and is thus ideally suited for the realization of such tectons. DNA origami utilizes large sets of oligonucleotides with rationally designed sequences to programmable fold a long single-stranded DNA scaffold into the desired shape. A range of well-behaved DNA origami barrels had been previously designed by Wickham et al. [33], with diameters ranging from 30 to 90 nm. In their approach, the outer barrel faces were used as a molecular canvas, where evenly spaced ssDNA overhangs were used as pixels to which other functionalities could be attached. For the design of our base monomer, we adopted the smallest reported barrel design and further reduced its height to 18 nm (as determined by TEM) to accommodate a shorter scaffold strand. This adjusted design requires fewer staple strands, which offers benefits for rapid and cost-effective prototyping.

We then reprogrammed the ssDNA overhangs to act as monomer connecting binding strands. As described in Figure 2a&b, the base monomer used in this work has 30 binding sites from which the binding strands originate. The sketch of the monomer structure in Figure 2a shows their distribution in alternating groups of 2 and 3 along 12 evenly distributed vertical binding stripes (2 × 6+3 × 6 = 30). The six binding stripes of each group of binding sites are arranged in six-fold symmetry, resulting in two overlaid principal six-fold binding site symmetries (Figure 2a).

**Fig. 2.**
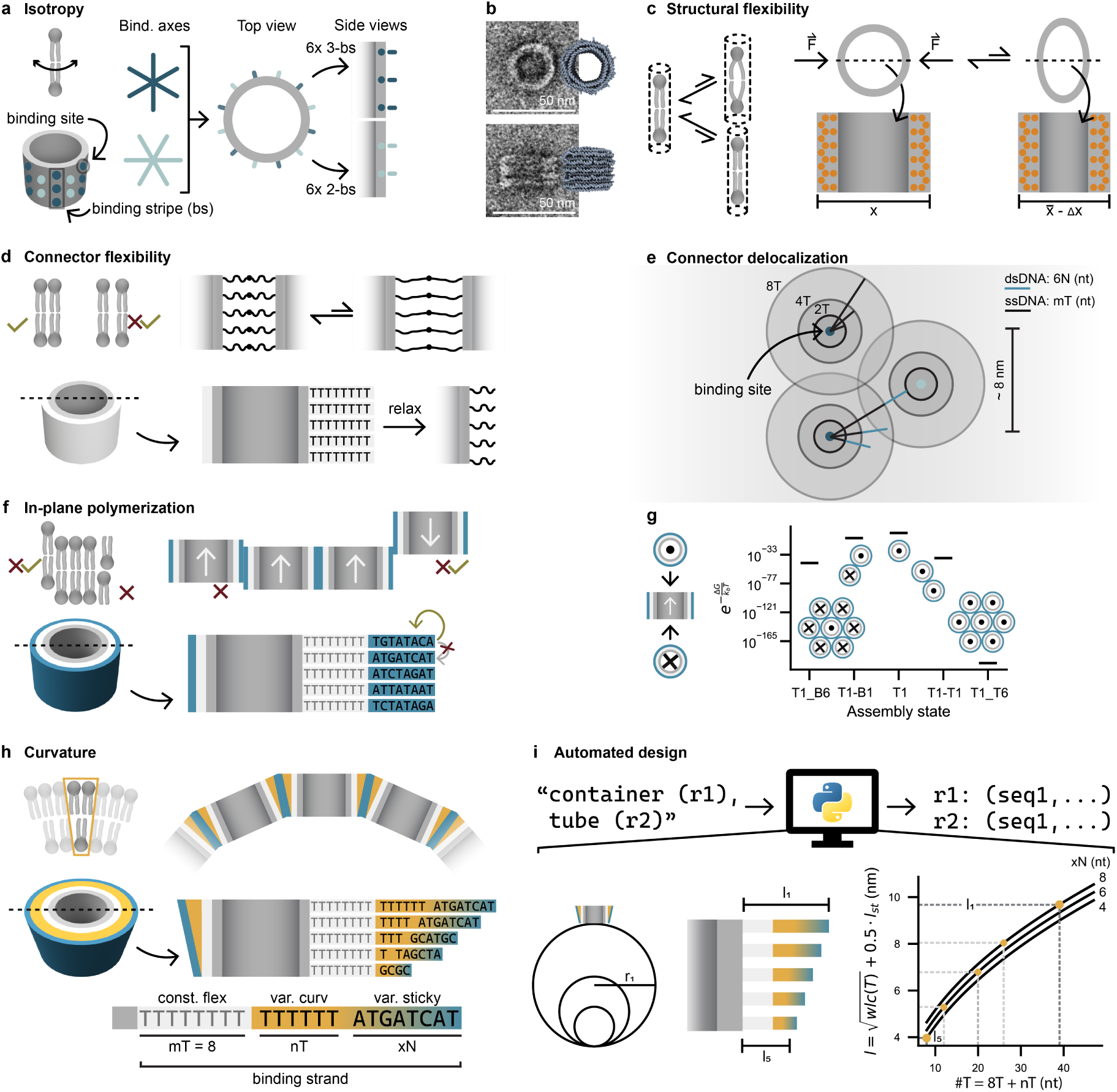
Principles for lipid-inspired monomer design. **a** Schematic of a DNA origami barrel and its binding sites. Thirty binding sites are distributed on the surface of the barrel in pairs of three (dark blue spots) or two (light blue spots), termed binding stripes (bs). Both binding site groups have a six-fold symmetry, with a relative rotation of 30° to each other. **b** Representative negative stain TEM micrographs of the DNA origami base-monomer, annotated with OxDNA simulated structures shown in the corresponding orientation. **c** Schematic indicating the compression of a flexible monomer under applied force, facilitated by its thin walls. Orange circles represent the positions of dsDNA helices. **d** Illustration of compliant monomer-monomer connections, achieved with flexible oligo dT domains in all ssDNA binding strands. **e** Visualization of the interaction cross-section for binding strands on the origami surface. Binding sites, denoting the origins of the binding strands, are shown as filled, dark blue or light blue dots for the two binding strip groups. The shaded areas represent the maximal ‘delocalization’ for flex domains with 2, 4, or 8Ts. Teal linker extensions indicate the contour length of the minimal binding strands with 6 nt sticky sequences, using inter-base distances of 0.63 nm for ssDNA [28] and 0.34 nm for dsDNA. **f** Planar growth of monomers in a specific binding orientation is enforced using a set of self-complementary, but otherwise orthogonal sticky domains (blue background). **g** The preference for planar growth can be rationalized via the Boltzmann weighted binding energies of a selection of sterically allowed assembly states. T1 represents a monomer oriented upward, B1 downward, and T1-B6 denotes a configuration of one upward-facing monomer encircled by six downward-facing ones. **h** Global curvature can be generated using asymmetric linker extensions with or without varying the lengths of the sticky domains (orange-blue). **i** Utilizing a Python script, we determine the inter-monomer angle that is required to assemble into a closed structure with a given curvature. Employing the worm-like chain (WLC) model, the script then generates binding strand sequences whose length matches the corresponding inter-monomer distances. Sets of tube or container forming binding sequences can be exported as CSV files.

Next to isotropy, we also sought to mimic the structural flexibility of lipids that is due in part to their aliphatic tails (Figure 2c). We thus deliberately used a barrel design with comparatively thin walls to provide some compliance, which allows the accommodation of defects and geometric mismatches through monomer deformation into flattened barrels or buckled hexagonal geometries.

In addition, all binding strands in one design begin with a constant length oligo-dT flex domain, making the connections more flexible (Figure 2d), increasing assembly error tolerance by preventing strain accumulation, and relaxing the requirements for the precise positioning of the binding sites. With longer flex domains, the binding sites are thus increasingly delocalized, resulting in a smeared-out, semi-continuous radial interaction profile, as sketched in Figure 2e. Notably, such effectively isotropic interactions are challenging to realize with rigid triangular or rectangular prisms as building blocks, which had been previously adopted for the assembly of other types of origami superstructures [18, 22]

Our basic DNA origami tectons can be readily polymerized to generate extended 2D membranes. As explained in Figure 2f&g, the specific design of the binding strand’s sticky domain enforces polymerization in the plane and discourages assembly in multi-layers or incorporation of monomers with the wrong orientation. Further, the use of multiple weak sticky domains combined with the Dipid binding site geometry should lead to a low nucleation probability, improving assembly formation [34].

For the monomers to assemble into curved structures, we focused on introducing local curvature through the monomer-monomer interactions, rather than explicitly designing pentamer subunit formation such as used for rigid, capsid-mimicking assemblies. Two monomers can be connected with up to five binding strands, defined by the binding stripes with either two or three ssDNA binding sites. Specifically, curvature can be introduced by asymmetrically extending the flex domains used for all binding strands with variable length curvature domains, composed of additional thymidines (Figure 2h). The angle *α* at which two neighboring monomers connect can thus be rationally controlled by varying the curvature domain and sticky domain lengths of specific binding strands.

In practice, we first created a simple geometric model of monomers bound to their neighbors at an angle *α* (Figure 2i), from which we obtained a set of five equally distributed inter-monomer distances corresponding to this angle. We next adopted the worm-like chain (WLC) polymer model to estimate the physical lengths of accessible binding strands with constant flex domains (mT = 8T), variable curvature domain lengths (nT = {0 − 52}T) and variable sticky domain lengths (xN = {4, 6, 8}T) (Figure 2i). With the used boundary conditions, an automated fitting procedure then generated a list of 74 unique sets of binding strands, which were each predicted to assemble into curved structures with a different target radius of curvature *r*.

## Self-assembly of small vesicle-sized containers

We expected monomers with isotropic curvature - i.e., monomers connected to all neighbors with the same angle *α* - to assemble into closed spherical containers (Figure 3a). Throughout this study, we arbitrarily selected and experimentally tested 6 container designs (XS, S, M, L, XL, XXL) (Figure 3b). In preliminary tests, we compared the appearance of structures that were based on the extra small vesicle design (XS), but had different flex domain lengths in their binding strands (mT = 2T, 4T, or, 8T). Transmission electron microscopy (TEM) images of these structures indicated that all designs formed spherical containers as desired (Figure 3c&d). As described in Figure 3c, when deposited on a TEM grid the containers tended to burst, but support by heavy staining or stabilization by neighboring containers can prevent their collapse, allowing more native structural imaging. Among the tested XS containers, the 8T flex-domain version appeared to be structurally more intact, and we thus decided to use 8T flex domains for all subsequent designs. Notably, all XS monomers formed containers directly after origami folding and without further purification.

**Fig. 3.**
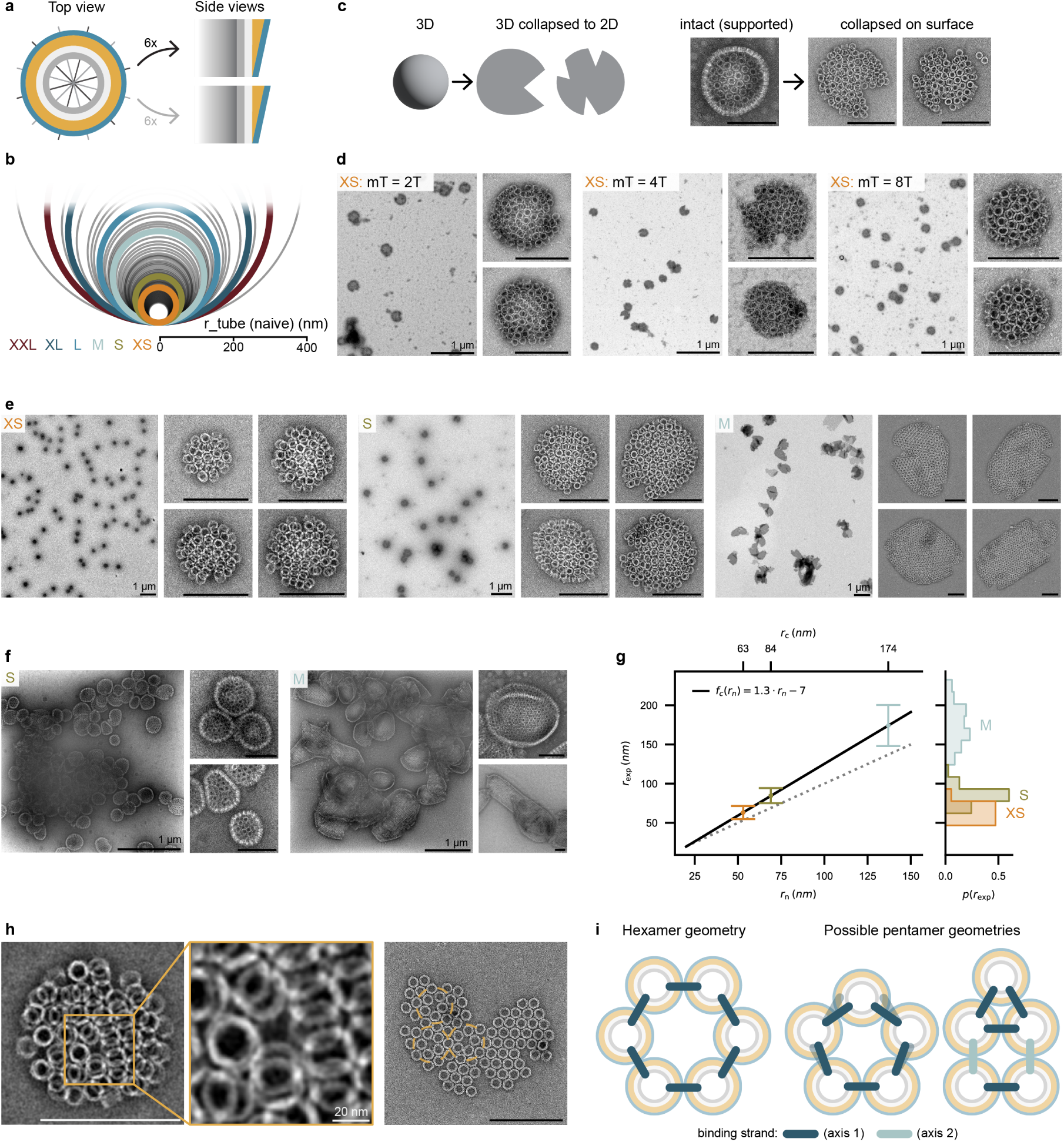
Dipid variants assembling into small and medium-sized containers. **a** Container-forming Dipids are designed to bind to all their neighbors at the same angle to generate a constant curvature, utilizing length variations in both binding stripe groups. Light and dark gray axes mark the binding stripe positions. **b** Predicted radii for all unique computed binding sequence sets. Our experimentally tested designs (XS, S, M, L, XL, XXL) are shown as thick, colored circles. **c** Illustration of the expected artifacts when 3D spheres break and spread in 2D, as it occurs during TEM grid preparation. On the right, example TEM images of an intact container supported by excess stain, as well as collapsed containers are shown. **d** Representative TEM overviews and detailed examples of XS containers formed from Dipid variants with a flex-domain length of m = 2, 4, or 8T. **e** Representative TEM overviews and detailed examples of containers formed from the XS, S, and M Dipid design variants. **f** Selected TEM overviews and detailed images of container aggregates. Mutual stabilization of the containers and excess staining helps to preserve their presumed native 3D shape. **g** Distribution of container radii, derived from TEM micrographs of flattened containers. Radii predicted by the naive model *r_n_* and the calibrated container model *r_c_* are displayed on the lower and upper x-axis, respectively. The black line is a weighted mean linear fit (error bars indicate standard deviation). Distributions of the experimental data are visualized as histograms on the right y-axis. **h** Example TEM micrographs showing monomers locally arranged into pentamers in collapsed and ruptured containers. **i** Schematic of a hexamer binding geometry, where all neighbor connections are achieved with binding sites from one group. Pentameric structures can form with the same binding strands as they are designed flexible (indicated with translucent strands). Alternatively, binding strands of both binding site groups can be combined to form pentamers. All scale bar lengths are set to 200 nm, if not specified otherwise.

Our design framework is highly modular. Changing the radius of curvature from the XS to the larger S and M designs (Figure 3b) was afforded by the exchange of only 24 DNA staple strands for each design. Whereas the XS and S monomers formed the targeted spherical containers at a high yield, our M design also formed closed structures, but these displayed a broader range of non-spherical morphologies (Figure 3e). This increase in morphological variability becomes even more apparent when comparing TEM micrographs of container aggregates (Figure 3f).

We next analyzed the principal radii of the XS, S, and M assemblies and compared them to the prediction of the naive model used for the initial design (Figure 3g). We then used our experimental data to refine the naive model, resulting in a calibrated model tailored to containers that was adopted for further designs.

Analyzing the structure of our assemblies in more detail shows that the barrels predominantly arrange into locally hexagonal geometries, but occasionally also form pentagons (Figure 3h). These are not, however, regularly spaced as in quasi-icosahedral (CK-like) assemblies, but appear at irregular locations. The formation of pentagonal assemblies is required for the formation of closed spherical shells and is, in fact, facilitated by the flexible and isotropic design of our origami monomers.

As the ssDNA binding strands are arranged in two groups with six-fold symmetry each, a pentagonal arrangement can be approximately achieved, by different arrangements of binding sites of both binding groups (Figure 3i).

## Self-assembly of cell-sized containers

Using our automated design approach, we next attempted to create even larger compartments (L, XL, XXL) that were predicted to exceed the size of the largest DNA origami containers realized so far. As judged from negative stain TEM micrographs, all three designs appeared to form closed structures, although these tended to burst when adsorbed to the substrate (Figure 4a). As the XL and XXL designs already reached sizes of 1 *µ*m and above, we were able to obtain additional evidence for the formation of closed containers by imaging them with conventional diffraction-limited fluorescence microscopy (Figure 4b). Compared to the predominantly spherical structures formed by small container designs, we increasingly observed elongated tubular or cone-shaped assemblies next to spherical vesicular structures. Figure 4c shows examples of the morphologies observed, which were all formed under the same conditions and using the same base monomer.

**Fig. 4.**
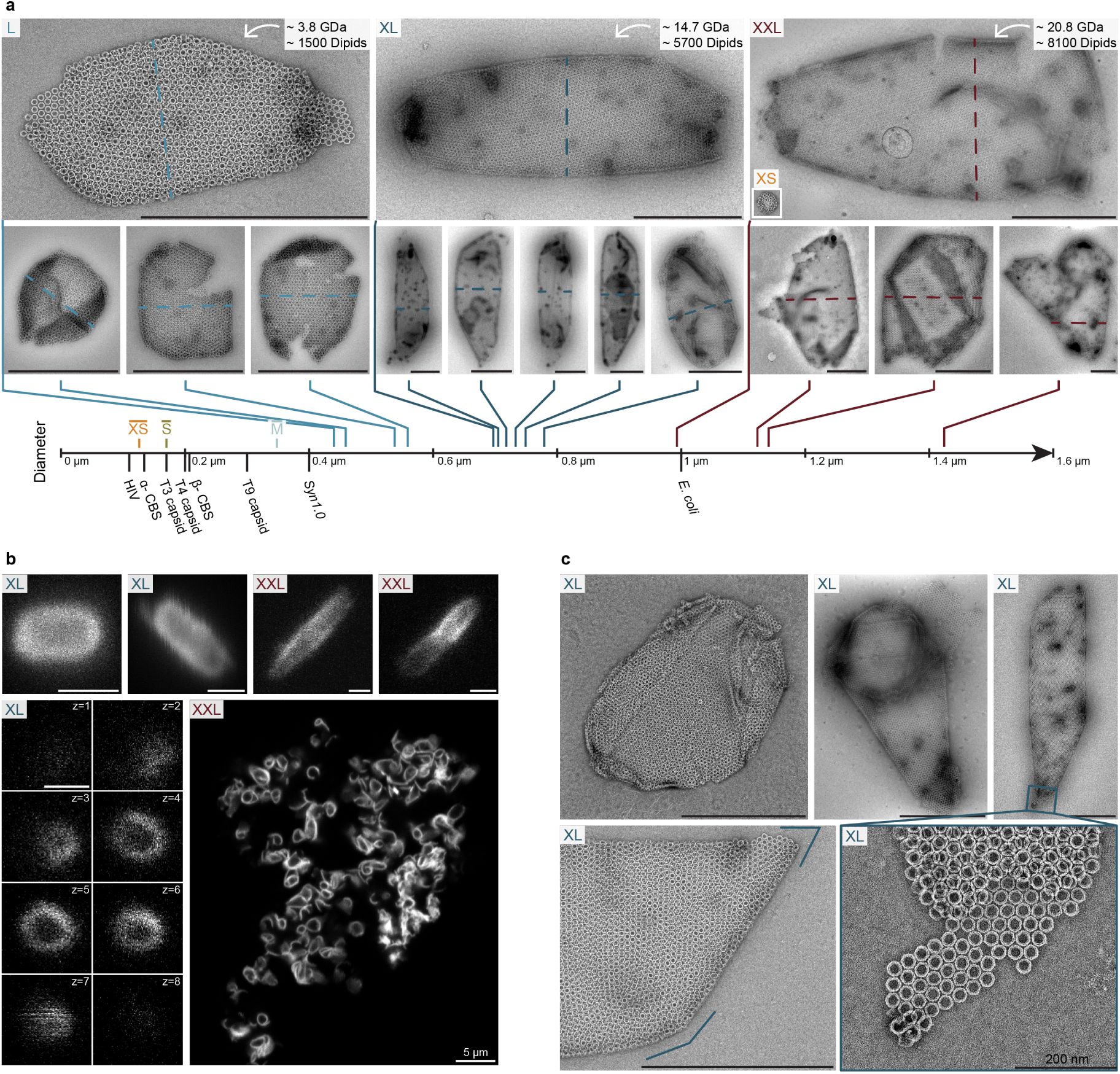
Giant containers formed by L, XL, and XXL Dipid designs. **a** Example TEM micrographs of isolated containers formed by the L, XL, and XXL designs. The inset image of the XS container in the large XXL example is displayed at the same pixel size. Container diameters were measured along the dotted lines shown in each image and back-calculated to their 3D diameters. Mean experimental diameters of XS, S, and M container designs are shown on the axis below. For comparison, dimensions of selected biological and state-of-the-art synthetic systems are also listed (dimensions were extracted from Refs. [18] (DNA origami capsids), [5] (HIV), [35] (*α*-Carboxysome : *α*-CBS), [36] (*β*-Carboxysome : *β*-CBS), and [37] (Syn1.0)). Scale bars: 1 *µ*m. **b** Laser scanning confocal images of XL and XXL assemblies. The close-up images in the first row are mean z-stack projections, while the lower left column displays eight slices of a z-stack, highlighting the hollow structure of the container. The right overview image shows a z-slice of an aggregate of assemblies containing partially open structures. **c** TEM micrographs depicting vesicle-like XL assemblies which, from left to right, increasingly display tubular morphologies. All containers are formed from the same monomers and imaged on the same TEM grid. In the lower part, examples of sharp corners as well as a close-up of an opened container end are shown. In the close-up, local pentameric assemblies are visible, as expected for fullerene-like cages. Scale bars: 1 *µ*m, unless specified otherwise.

The increasing morphological variability, ranging from approximately spherical containers (XS, S) to fullerene-like containers (M, L), and finally to closed tube-like containers (XL, XXL), can be readily explained by existing models [2, 38, 39]. Using continuum elasticity theory and simulations, four morphological assembly regimes were identified for flexible, triangular monomer assemblies [39]. According to these, the dominant morphology of structures formed from a single monomer type depends on both the designed radii and the membrane’s elasticity. Apart from the three morphological regimes we observed experimentally, theory predicts that further increasing the designed container radii will eventually lead to the formation of open tubes, once pentamer formation becomes highly unlikely. We anticipate that the morphology of containers with a designed size can be influenced by adapting the membrane elasticity through the lengths of the oligo dT flex-domains, or by modifying the energetic barrier of pentamer formation. Another approach to controlling morphology could involve tuning the container assembly conditions, inspired by the commonly observed morphological variability in virus particles and their strong dependence on assembly conditions [4, 40–42].

## Tube formation with controlled mean radii

Intrigued by the tendency of the larger designs (XL, XXL) to form tubular assemblies, we decided to adapt their monomer design to deliberately promote the formation of open tubes with controllable radii. The formation of open tubes does not require any of the monomers to arrange into pentagons, which allowed us to remove all binding stripes of one of the six-fold binding axes and thus reduce the number of binding strands from 30 to 18.

Figure 5a shows how we implemented the two principal tube curvatures 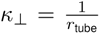 and *κ_||_* = 0 with the simplified tube Dipid. First, we designed three sticky domain sets to enable specific binding stripe interactions. As shown in Figure 5a, the binding strands of two opposing binding stripes are always chosen the same, resulting in an overall two-fold symmetry.

**Fig. 5.**
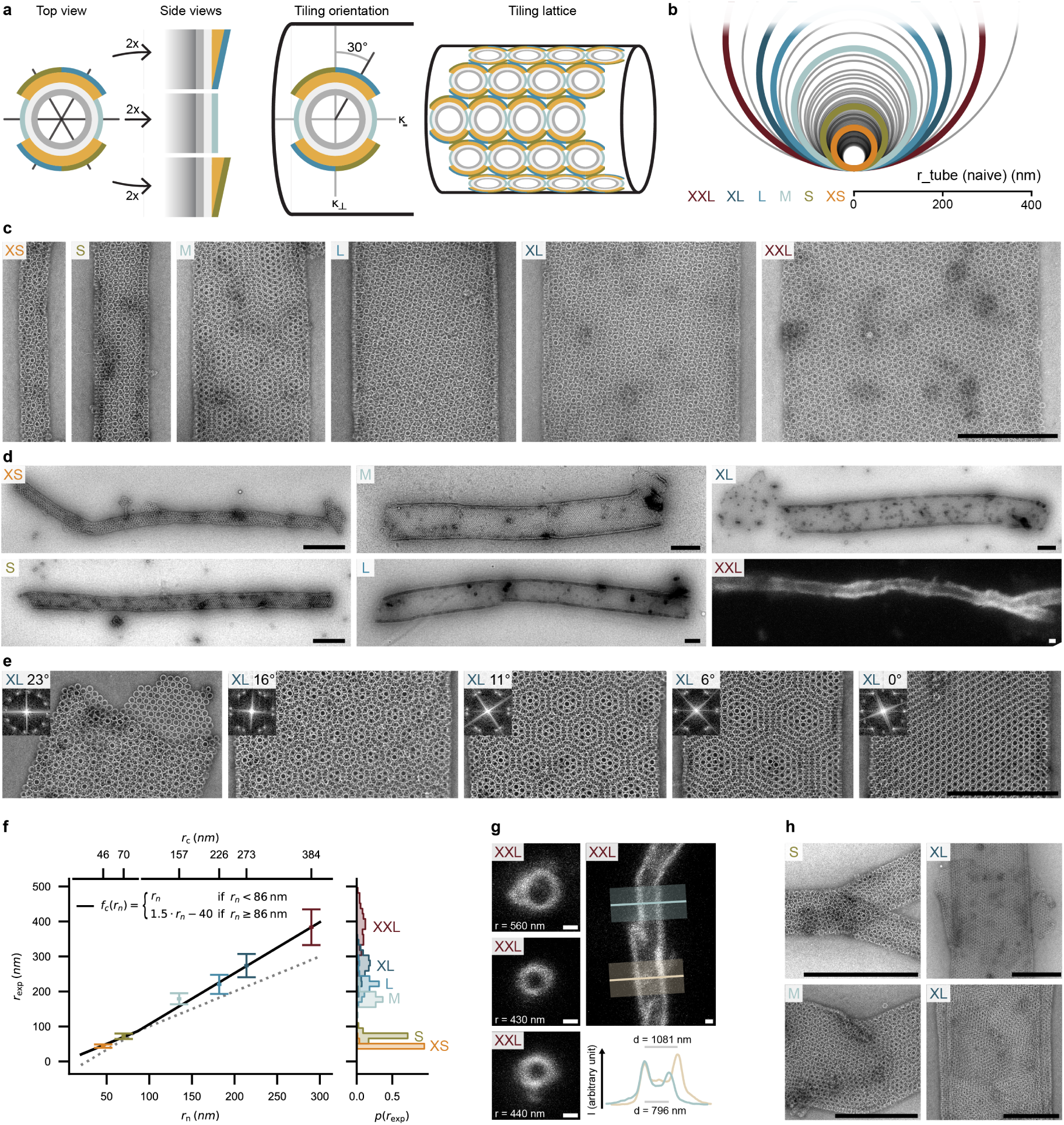
Formation of open tubes with programmable radii. **a** Monomer design for the formation of tubes, for which only binding stripes from one set of binding axes are utilized (cf. Figure 2a). Differently colored sticky layers (green, blue, light blue) indicate binding strand sets with orthogonal sticky domains. Orange layers are introduced to induce curvature perpendicular to the tube axis as indicated in the tiling lattice. **b** Predicted tube radii for all computed design variants. Experimentally tested designs (XS, S, M, L, XL, XXL) are shown with thicker outlines and coloration. **c** TEM micrographs of tubes with incrementally increasing designed radii from left to right. All images are shown at the same pixel size. **d** Exemplary overview images showing the morphology of tubes at their full elongation. XS-XL tubes are shown as TEM micrographs, while the XXL-tube image is a confocal microscopy z-stack maximum projection. **e** Gallery of Moiré patterns acquired from a single sample of XL tubes imaged on the same TEM grid. Insets show fast Fourier transforms (FFT) of circular regions within the images, from which the relative angle between the upper and lower hexagonal grids formed by the tube walls can be derived. **f** Distribution of tube radii, back-calculated from the diameters of flattened tubes observed via TEM. Since radii can vary along the length of a tube, several measurements were taken for each tube at distances of ≈ 1.5× the tube diameter. Radii predicted by the naive model *r_n_* and the calibrated tube model *r_c_* are displayed on the lower and upper x-axis, respectively. The black line is a weighted mean fit to the experimental radii. Error bars indicate the standard deviation. Distributions of the experimental radii are shown as histograms on the right. **g** Close-up confocal microscopy maximum-value projections. In solution, tubes can be oriented orthogonal to the imaging plane, displaying their cross-sections (left column), and parallel to the plane, illustrating the variation in cross-section along a tube’s length (right column). **h** Selected TEM micrographs showing commonly encountered defects such as branching, bending coupled with diameter changes, and growth of ‘skirts’ or multi-layered tubes. Scale bars: 500 nm, unless specified otherwise.

*κ_||_* = 0 is implemented with an equal-length binding strand set, generating zero curvature along that polymerization axis. The binding strand sets of the remaining binding stripes differ only in their sticky domains and are chosen to generate the desired tube curvatures (Figure 5a&b).

As the curvature generating binding stripes are oriented at 30°C with respect to the tube’s diameter, the resulting tube radius is *r*_tube_ = *r*_monomer_ ·cos (30°C). Using this correction, we can use our original list of binding strands to design tubes for 62 unique targeted *r*_tube_. As the binding strands implementing *κ_||_* are identical for all tubes, as are the shortest binding strands of each curvature inducing binding stripe, changing from one tube radius to another requires the exchange of only up to 8 staples.

We chose six computationally generated designs with different predicted tube diameters and imaged the resulting assemblies using negative stain TEM and confocal microscopy (Figure 5c&d). All tested designs assembled as desired into tubular structures. Notably, we rarely observed smaller, misassembled structures or sheet-like assemblies that had failed to form tubes. This suggests that tube growth is considerably faster than the formation of new seeds. Occasionally observed aggregation or interconnection of several tubes (SI FigureA1) might be due to the incorporation of rarely occurring origami dimers, or due to salt-mediated electrostatic attraction between tubes.

For all of our tube assemblies, we consistently observed intriguing Moiré patterns in the TEM micrographs (Figure 5c&e), which result from the superposition of the hexagonal origami lattices on the front and the back of the tubes. Depending on the relative orientation between these lattices, different patterns emerge. The rotation angle can be inferred directly from the pattern, or measured using Fourier transforms of the images (Figure 5e).

We next compared the radii of the tubes determined by TEM and confocal microscopy with those predicted by our naive model. With a mean absolute error of only 2 nm, the prediction of the naive model for the two smallest tubes turned out to be surprisingly accurate. The larger tubes were found to be larger than anticipated, however, showing close to linear deviation from the naive prediction (Figure 5f). We believe that the observed discrepancy results from our application of a polymer mechanical model for the ssDNA binding strand lengths in a regime where it is actually not expected to be accurate. Specifically, the binding strands used for the larger radii contain only short ssDNA domains (up to 10 nt), which are less well described as polymers than the comparatively longer ssDNA domains employed for the smaller radii (up to 37 nt). In order to improve the prediction of tube radii for future designs, we heuristically fitted lines to the data in the two regimes. The resulting calibrated model tailored to tube designs has a mean absolute fitting error of only 5.5 nm, making it sufficiently accurate for practical use.

As expected [39, 43], the distribution of the radii of tubes formed from a single Dipid variant becomes broader with increasing tube radius (Figure 5f). In the initial growth phase, the monomers are expected to assemble into a curved sheet, which at one point will close back on itself to form a tube seed that will continue to grow into a longer tube of the same diameter. The tube closure event is expected to occur statistically, with larger fluctuations for the larger tube sizes. Furthermore, the tubes may form with different chiralities, depending on whether the tube is closed precisely along the direction defined by the curvature of the monomers, or with a slight shift. The shifted tube-closure events give rise to the observed Moiré patterns discussed above. If precise tube radii are required, one could increase the assembly complexity at the cost of assembly time [44].

Notably, we occasionally observed the same tube to display different Moiré patterns at different locations, suggesting a variation of tube diameter along the tube. We encountered other growth irregularities such as branching, bending, and the growth of structures from defects, potentially leading to multi-layer tubes (Figure 5h).

## Conclusion

With the Dipid framework, we introduce a lipid-inspired approach to generate large super-assemblies from DNA origami subunits, capable of forming extended 2D layers as well as closed compartments with controllable sizes ranging from nanometers to micrometers. With 25 Gigadalton, our largest compartments are the largest demonstrated with the tools of DNA nanotechnology so far.

Our framework represents a significant departure from the traditional approach in the field, which previously focused on the realization of super-assemblies from precisely defined, structurally rigid monomeric origami subunits. Using only a single, highly symmetric origami monomer unit with flexible and redundant binding interactions, the realization of superstructures is comparatively simple, easily adaptable, and cost-effective, and thus has the potential to be widely adopted for the realization of DNA super-assemblies. Notably, based on the modular design approach outlined here, one can modify existing and develop novel Dipid based compartments without extensive previous experience in DNA origami design.

As demonstrated above, our larger super-assemblies easily reach the size of small unicellular organisms, or those of large unilamellar vesicles made from phospholipids. As such, they represent interesting alternatives for the compartmentalization of synthetic biological systems in bottom-up biology. Compared to vesicles, among the potential benefits of Dipid compartments are their designable sizes and curvatures, as well as their structural stability combined with the porosity of the monomers.

As the interactions between the monomers are only encoded within a small subset of the DNA origami staples, most of them remain available for modification, as does the Dipids interior. Hence, compartments could be formed from monomers that carry different chemical or mechanical functions, e.g., enzymes, nanomachines, or inorganic components. Integration of DNA-based modules for sensing and signaling [45], computation [46], bioproduction [47], or locomotion would thus allow to program the compartments as soft cell-scale robotic systems.

One of the challenges for applications as cell mimics, small-scale robots, or bioreactors will be the encapsulation and sequestration of functional components inside of the compartments, potentially supported by further sub-compartmentalization with synthetic organelles. Supplying such systems with nutrients or substrates could be readily achieved through the large pores of the Dipid membranes.

In principle, it should be possible to assemble similar compartments using RNA-based monomers [48–51], which would open up the possibility to create nucleic acid-based protocells, whose contents and boundaries are both generated by a transcription process.

## Methods

### Design of Binding Domains Using NUPACK

The orthogonality of sets of binding domains was evaluated utilizing the NUPACK web service (Model: dna04, Temperature: 25 °CC, Maximum complex size: 2, Strand concentration: 1 M). The hybridization free energies were computed with the NUPACK Python package [52] (material=“dna”, celsius=20 °CC, sodium=0.05 M, magnesium=0.018 M).

### Binding Strand Length Calculations for Specific Curvatures

We determined the binding strand lengths by maintaining a constant flex domain length (mT = 8T), varying the sticky domain lengths (xN = 4, 6, or 8N), and altering the curvature domain lengths (nT = 0 − 52T). We calculated the contour lengths of ssDNA domains *L_c_*(ssDNA) and dsDNA domains *L_c_*(dsDNA) were calculated using a base-to-base distance of 0.63 nm base^−1^ for ssDNA [28] and 0.34 nm base^−1^ for dsDNA. The persistence length of ssDNA, *L_p_*, was assumed to be 1.5 nm [28].

Binding strand lengths *l*_strand_ were calculated using the equation

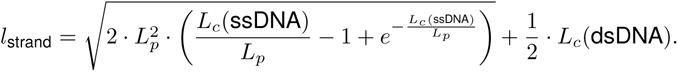

We then employed a simplistic geometric monomer model to identify sets of binding strand lengths that would enable the monomers to bind at a designated angle. The monomer model dimensions, derived from TEM analyses, were defined as a width of 28.5 nm and a height of 18 nm. Subsequently, we assigned appropriate binding strand sequences to each necessary linker length. Initial naive radii predictions were refined using container and tube calibration functions.

A Python script for this procedure is available at our GitHub repository.

### Calculation of free energies for different monomer arrangements

We estimated the Δ*G* for each of the assemblies shown in Fig. 2g by multiplying an average linker hybridization energy of 42.85 kJ mol^−1^ by the count of sterically feasible bonds between a central monomer in upward orientation and its nearest neighbors. We assume that monomers are arranged to bind predominantly through the 3-binding stripes. In the T1-T6 configuration, where 2×3 compatible binding strands are positioned between three monomers, only two of each type can hybridize simultaneously. Thus, we considered these connections as 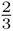 of a full bond. The analysis was conducted at a temperature of 20 °CC.

### OxDNA Simulations

DNA origami structures were exported from scadnano [53] and pre-arranged using oxView [54]. The forces used during structure relaxation were also generated in oxView. After a short Monte Carlo pre-relaxation step, we employed Molecular Dynamics (MD) simulations (with the oxDNA2 model [55]) to first relax and then simulate the monomer structure. We generated PDB files for rendering in ChimeraX with the oxDNA analysis tools command line script [55]. The monomer structure presented in this article represents a mean structure from an equilibrated simulation. The Python code used for setting up and running the relaxation and simulation runs, along with all input and parameter files, is available on our GitHub.

### DNA Origami Design and Folding

We adapted the DNA origami structures developed by Wickham et al. [33] using scadnano [53]. Structures were folded with a p2873 scaffold (provided by Prof. Hendrik Dietz’s group, 100 nM in ddH2O) and varying sets of staple strand oligonucleotides (Integrated DNA Technologies, 200 µM in 20 mM TRIS, 0.1 mM EDTA, pH 8.0). Folding reactions were prepared with a final scaffold concentration of 50 nM and a staple strand concentration of 200 nM in FOB18 buffer (5 mM TRIS, 1 mM EDTA, 18 mM MgCl_2_, and 5 mM NaCl). An annotated scadnano file and the scaffold sequence are available on our GitHub. Folding solutions were annealed in a thermocycler (Eppendorf, Mastercycler nexus GX2) using different annealing ramps for different designs: for designs without sticky-strand extensions, 15 min at 60 °CC followed by a decrease of 0.1 °CC every 6 min from 56 °CC to 53 °CC; for containers and tubes, 15 min at 70 °CC followed by a decrease of 0.1 °CC every 18 min from 65 °CC to 20 °CC. We stored samples at room temperature for subsequent use.

### DNA Origami Purification

Samples were purified using ultrafiltration with FOB5 washing buffer (5 mM TRIS, 1 mM EDTA, 5 mM MgCl_2_, and 5 mM NaCl). A centrifuge (Eppendorf, Centrifuge 5425 R) was first pre-heated to 32 °CC. For each folding solution, we loaded 1.5 mL of FOB5 buffer and centrifuged for 15 min at 10 krcf and 32 °CC. We then placed Amicon Ultra 0.5 ml Ultracel filters (100k) in the centrifuge, loaded with 500 µL of 32 °CC FOB5 buffer, and centrifuged at 10 krcf for 5 min. We discarded the supernatant. Next, we loaded the filters with 430 µL of 32 °CC FOB5 buffer and 60 µL of folding solution, and centrifuged at 10 krcf for 5 min, discarding the supernatant afterward. This step was repeated with 450 µL of 32 °CC FOB5 buffer, followed by centrifugation at 10 krcf for 5 min, and again the supernatant was discarded. Finally, we loaded 450 µL of FOB18 assembly buffer at room temperature and centrifuged at 10 krcf for 5 min. We extracted the purified and buffer-adjusted samples from the filters with a pipette by repeatedly aspirating and dispensing the solution to dissolve any potential pellets.

### Optional Re-Assembly After Purification

After purification and buffer adjustment to FOB18, we re-assembled the DNA origami samples (tubes and containers already form during the folding process) in a thermocycler (Eppendorf, Mastercycler nexus GX2). We disassembled the structures at 38 °CC for 30 min, followed by an annealing ramp of −0.1 °CC every 12 min from 38 °CC to 20 °CC.

### Acquisition of TEM Micrographs

For TEM imaging, we incubated 5 µL of DNA origami samples on glow-discharged (coating time:20s, coating current: 35mA, polarity: negative) formvar carbon Cu400 TEM grids (Science Services) for 30-300 s, depending on DNA origami concentration. We prepared the staining solution by adding 1 µL of 5 M NaOH to 200 µL of a 2% uranyl formate solution, followed by vortexing and subsequent centrifugation at 5 °CC and 21 krcf for 5 min. After incubating the samples, we washed the grids with 5 µL of stain and then incubated them with 15 µL of stain for 30 s. Imaging was conducted with a FEI Tecnai T12 microscope (120 kV) equipped with a Tietz TEMCAM-F416 camera, operated with SerialEM.

### Post-Processing of TEM Micrographs

We applied bandpass filter cutoffs between 1-3 px for the lower boundary and 300-1000 px for the upper boundary. Micrographs were locally contrast-adjusted to improve structure visibility, with or without prior bandpass filtering. For the Enhance Local Contrast (CLAHE) function in Fiji, we used block sizes between 32 and 127 px, a maximum slope of 3 or 4, and histogram bins set to 512.

### Radii Extraction from TEM Micrographs

We recorded overview TEM micrographs for each design and manually measured diameters with Fiji. For containers, measurements were taken along their narrow axis if aspect ratios deviated from unity. Tube measurements followed two guidelines:

1. Diameters of the same tube were measured with a minimum distance of twice the average tube diameter.
2. Diameters were measured only at positions where the local tube diameter appeared uniform over a width of at least one tube diameter.

Extracted diameters *d*_2D_ were used to calculate the radius of the assumed 3D structure *r*_3D_ with 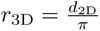. Structure diameters mentioned in this work refer to *d*_3D_ = 2 · *r*_3D_.

### Light Microscopy

Each DNA origami monomer was designed to include one Atto655 dye labeled DNA oligonucleotide (Biomers), protruding into the inside of the barrel. We imaged DNA origami samples in passivated ibidi *µ*-slides VI 0.1, using either a Nikon Ti-2E microscope (NIS elements software, SOLA SM II LED light source, Andor NEO 5.5 camera) or, for higher-resolution images, a Leica TCP SP8 laser scanning confocal microscope (LAS X software).

### Light Microscopy Image Post-Processing

We globally adjusted the contrast of light microscopy images using Fiji [56]. Z-stack projections, either maximum or mean, were also created using Fiji.

### Moiré Pattern Analysis

Tube sections with prominent Moiré patterns were selected from the same TEM grid of a purified and annealed XL tube sample. FFTs were computed from unprocessed TEM micrographs using a circular region of interest. For visualization, images were bandpass filtered (3-300px) and globally contrast-adjusted. We determined the rotational angle of overlapping hexagonal grids in each image based on the innermost maxima of each FFT. Using Fiji’s “find maxima” tool, we detected the maxima and extracted angles relative to the image frame using Fiji’s “measuring” tool. We computed the mean angular shift and its standard deviation using a Python script (GitHub).

### Large Language Model Tools

We used the online tool ChatGPT [57] (GPT4 model [58], OpenAI) to assist with programming.

## Supplementary information

## Acknowledgments

This research was conducted within the Max Planck School Matter to Life, supported by the German Federal Ministry of Education and Research (BMBF) in collaboration with the Max Planck Society. We acknowledge support by the TUM Innovation Network “Robotic Intelligence in the Synthesis of Life (RISE)”, which is financed through the Excellence Strategy of the Federal Government and the Länder. The authors thank E. Kopperger and J. List for helpful discussions, S. Wickham for support and inspiring discussions on her barrel designs, A. Pastucha for confocal microscopy training, and M. Kube for TEM training. Correspondence and requests for materials should be addressed to F.C.S.

## Ethics declarations

### Availability of data and materials

All datasets are available upon request.

### Code availability

Python code used for sequence generation is publicly available at our GitHub repository at the URL “https://github.com/ckarfusehr/Cell_scale_DNA_origami_membranes”.

Appendix A Appendix

**Fig. A1.**
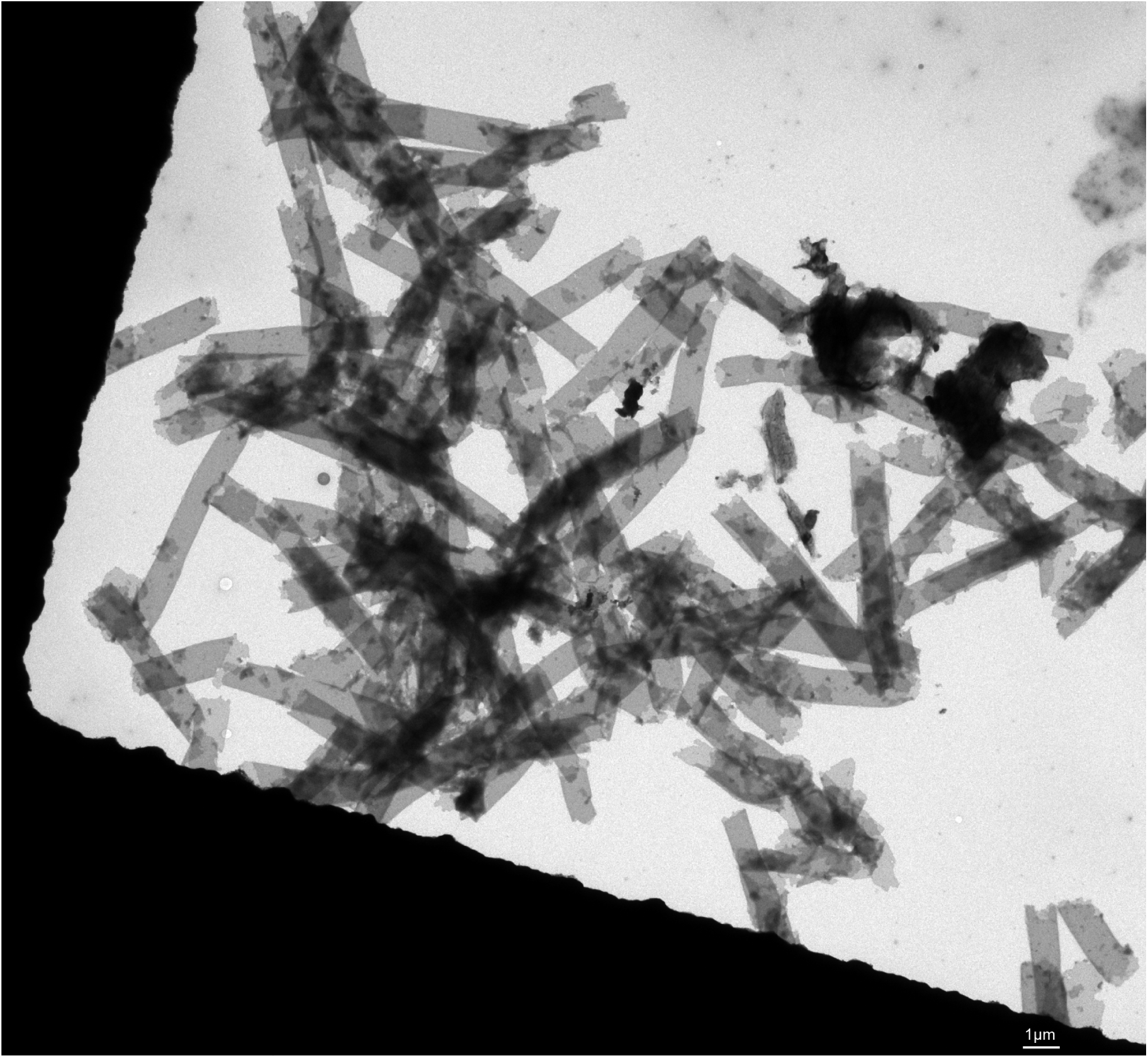
Example TEM micrograph of aggregated tubes, observed in a purified and re-annealed sample of the L tube variant.

